# Indexed variation graphs for efficient and accurate resistome profiling

**DOI:** 10.1101/270835

**Authors:** Will P.M. Rowe, Martyn D Winn

## Abstract

**Background:** Antimicrobial resistance remains a major threat to global health. Profiling the collective antimicrobial resistance genes within a metagenome (the “resistome”) facilitates greater understanding of antimicrobial resistance gene diversity and dynamics. In turn, this can allow for gene surveillance, individualised treatment of bacterial infections and more sustainable use of antimicrobials. However, resistome profiling can be complicated by high similarity between reference genes, as well as the sheer volume of sequencing data and the complexity of analysis workflows. We have developed an efficient and accurate method for resistome profiling that addresses these complications and improves upon currently available tools.

**Results:** Our method combines a variation graph representation of gene sets with an LSH Forest indexing scheme to allow for fast classification of metagenomic sequence reads using similarity-search queries. Subsequent hierarchical local alignment of classified reads against graph traversals enables accurate reconstruction of full-length gene sequences using a scoring scheme. We provide our implementation, GROOT, and show it to be both faster and more accurate than a current reference-dependent tool for resistome profiling. GROOT runs on a laptop and can process a typical 2 gigabyte metagenome in 2 minutes using a single CPU.

**Conclusion:** We present a method for resistome profiling that utilises a novel index and search strategy to accurately type resistance genes in metagenomic samples. The use of variation graphs yields several advantages over other methods using linear reference sequences. Our method is not restricted to resistome profiling and has the potential to improve current metagenomic workflows. The implementation is written in Go and is available at https://github.com/will-rowe/groot (MIT license).

## Background

Antimicrobial resistance (AMR) remains a significant global threat to public health, with directly attributable deaths per annum predicted to rise from 700,000 to 10,000,000 by the year 2050 [1]. This threat is not restricted to humans, with consequences also for animal health, welfare and food production [2]. In an effort to inform policy that can mitigate the spread of AMR, there has been a recent drive to establish surveillance programmes to monitor the prevalence of AMR [3, 4]. These programmes have traditionally used culture or PCR-based surveillance techniques, restricting monitoring to a few key genes or organisms. However, the use of metagenomics for surveilling AMR is gaining traction as it offers a much higher throughput over these traditional techniques [5, 6].

The use of metagenomics to determine the antibiotic resistance gene (ARG) content of a microbial community, hereafter referred to as resistome profiling, has been applied in studies of a wide variety of biomes [7–16]. Studies such as these utilise several well-maintained ARG databases [17–19] and ARG-annotation tools [20–23], of which only a few are designed for resistome profiling (Figure 1). These databases and tools are used with assembled metagenomic contigs, or alternatively with metagenomic sequence reads. In the case of the latter some tools and studies opt to align reads to reference ARGs and report only fully-covered references [10], whereas some bin reads using BLAST and similarity/length thresholds [16]. All of these strategies offer the user a compromise between ease of use, analysis speed and the accurate typing of ARGs (ability to detect ARG type/subtype)

**Figure 1.**
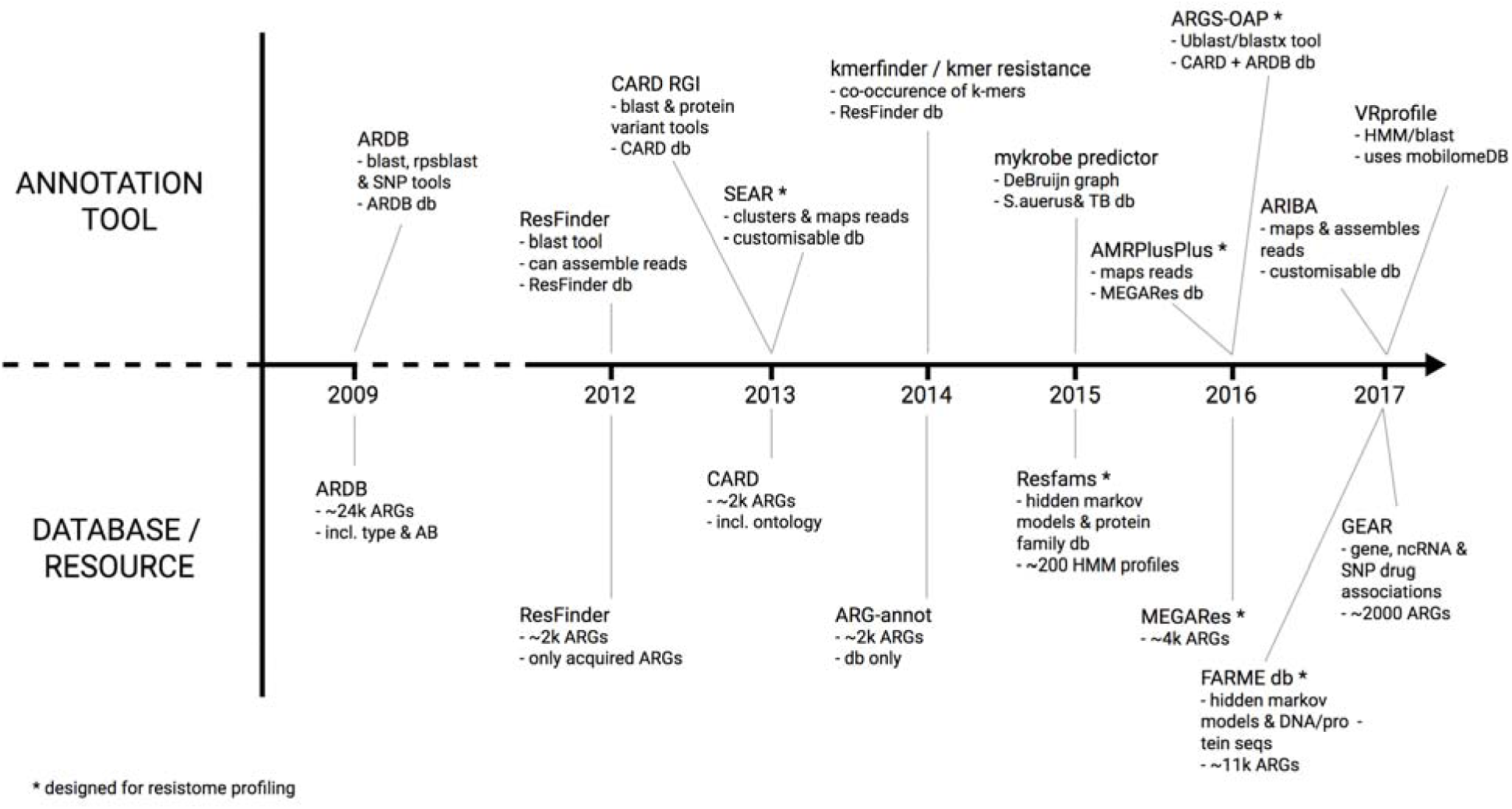
Summary of ARG annotation tools and databases. This figure shows several published tools and databases that are used to detect ARGs (in sequencing data or assembled contigs). The list is not exhaustive and the tools that are designed specifically for resistome profiling are highlighted.

One of the main limitations of existing tools for resistome profiling (or profiling of other gene sets) in metagenomes is that high similarity shared between reference sequences can result in ambiguous alignments, unaligned reads (false negatives), or mis-annotated reads (false positives); thus reducing the accuracy when typing ARGs [24]. A solution to this has been to cluster the reference sequences and then use the cluster representative sequences as a target for read alignment [20], however this results in a loss of information as ARG subtypes will be masked. Likewise, a related approach has been to collapse final annotations of ARG variants into a common ARG annotation, also masking subtypes [14]. The ability to accurately detect ARG type/subtype, here termed profiling resolution, is important considering that the variation between subtypes of ARGs can result in different phenotypic activity [25].

Recently non-linear data structures, such as variation graphs, have been used to encode reference sequences for applications such as variant calling [26, 27]. In terms of ARG annotation, the Mykrobe predictor tool applies non-linear reference representation in the form of a de Bruijn graph to identify resistance profiles in *Staphylococcus aureus* and *Mycobacterium tuberculosis* isolates [28]. Non-linear reference representation reduces redundancy whilst maintaining information that facilitates classification. Variation graphs, which are directed acyclical graphs, are offered as a solution to reference bias in population genomics as they represent sequence variation within a population [27]. In order to align a sequence against a variation graph, the traversals within a graph are indexed. Variation graph indexing approaches to date have included hash-map and Burrow-Wheeler transform encoding [29, 30]. Index design must balance query length, performance and index size in order to deal with the complexity of variation graphs. Efficient indexing is further complicated in the case of multiple graphs.

We propose that variation graph traversals can be indexed using Locality Sensitive Hashing (LSH), allowing for fast and approximate assignment of a sequence to a specific region of a traversal within one or more graphs. MinHash is a form of LSH that can be used to compress sets into smaller representations, called MinHash signatures. MinHash signatures can be used to estimate the similarity of the original sets independently of the original set size and was first used for duplicate webpage detection [13, 31]. The MinHash technique is increasingly being used in bioinformatics applications for clustering and searching large sequencing datasets [32, 33] and has also recently been applied to read alignment algorithms [34–36]. In applications such as these, the ability to efficiently compare signatures is essential. Additional LSH indexing techniques can therefore be applied to reduce dimensionality and facilitate querying of signatures. One such technique is the LSH Forest indexing scheme that uses multiple prefix trees to store hash tables containing portions of the MinHash signatures and facilitates self-tuning of index parameters, offering a performance improvement over traditional LSH indexes [37].

In this paper, we present a method for resistome profiling that utilises a variation graph representation of ARG databases to reduce ambiguous alignment of metagenomic sequence reads. We store sets of similar ARG reference sequences in variation graphs; collapsing identical sequences whilst retaining unique nodes that allow for accurate typing. By applying an LSH Forest indexing strategy to the variation graph collection that allows for fast and approximate search queries, we show that metagenomic reads can be seeded against candidate graph traversals. Subsequent hierarchical local alignment of reads and scoring enables accurate and efficient resistome profiling. We also provide our implementation, Graphing Resistance Out Of meTagenomes (GROOT), and compare it against ARGS-OAP, a recently published tool for resistome profiling [23].

## Methods

Here we describe our method to index a collection of ARG reference sequences, align sequence reads using the index and then apply this for resistome profiling of metagenomics samples. We then document our implementation, GROOT, and describe how we evaluated its performance.

### Indexing

The first stage in indexing a collection of ARG reference sequences is to remove redundancy, thus reducing the number of potential alignments to multiple references by collapsing identical sequence regions into a single reference. To do this we employ a Directed Acyclical Graph (variation graph) representation of ARG reference sequences. By fingerprinting variation graph traversals and using this as an index, we can then perform approximate seeding of sequence reads (Figure 2) (Algorithm 1 in Additional File 1).

**Figure 2.**
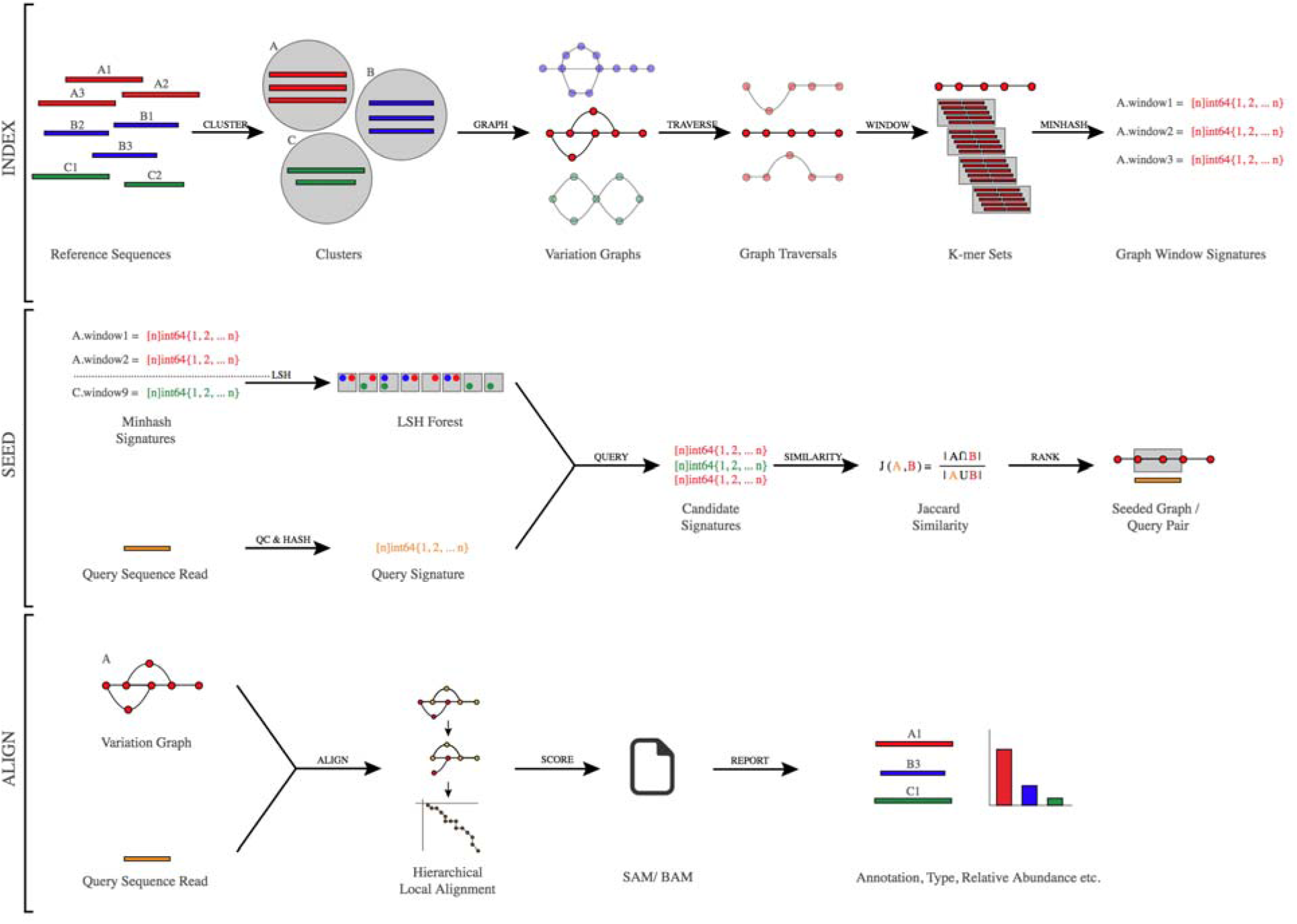
Overview of our method to index and query ARG variation graphs.

#### i. Create variation graphs

To convert a collection of FASTA-formatted ARG reference sequences to a set of variation graphs, all sequences are first clustered based on sequence identity (∼90%) in order to group sequences into distinct sets of similar sequences. Each resulting set of similar sequences can be viewed as a Multiple Sequence Alignment (MSA) that describes the insertions, deletions and polymorphisms in each member of the set, relative to a set representative sequence. The MSA is converted to a variation graph by first using the representative sequence as the graph backbone; each base of the sequence is a node with edges connecting them in series. The other sequences of the MSA are then incorporated; common bases are consolidated as single nodes and all deletions and insertions are added via edges and bubbles.

Each variation graph node contains information such as parent sequence, reference position and encoded base, allowing restriction of graph traversals to known references and the translation of traversals into reference name and sequence location. In the case of sets containing a single sequence, they are still converted to variation graphs but will be a linear series of nodes and only contain one possible traversal. All variation graphs are topologically sorted so that node ordering reflects MSA position prior to fingerprinting graph traversals.

#### ii. Fingerprint graph traversals

The second indexing stage is to fingerprint the traversals in each variation graph, allowing for fast and approximate matching of query reads to region(s) of a variation graph. To fingerprint a graph, a sliding window of length *w* is moved across all graph traversals and a MinHash signature is created for each window [31]. The windows are typically the same length as the expected query reads. The starting nodes of consecutive windows are spaced by an offset, *o*, where *o* ≤ *w*.

For each window, the contained nodes are joined to form a sequence (encoded as an array of bytes), which is then decomposed into a set of k-mers of length *k*, where *k* < *w*. To then create a MinHash signature (an array of unsigned 64-bit integers) of length *s*, each k-mer in the set is evaluated in series. A k-mer is hashed to an unsigned 64-bit integer *s* times, using *s* distinct hash functions. Each time a k-mer is hashed, the index position in the signature is advanced from 0 and the new hash value is evaluated against the value stored at the current index position; if the stored value is greater than the new value, the stored value is replaced by the new value. The signature index is reset to 0 prior to hashing the next k-mer in the series and the same *s* hash functions are used. Once all k-mers in a set for a given window have been hashed, the signature for that window is complete; the signature is linked to the graph ID and the window start node before it is stored. A single graph node can have multiple linked signatures if multiple traversals are possible.

#### iii. Store window signatures

The final indexing stage is to store the window signatures for each variation graph in a data structure that allows for fast and approximate nearest-neighbor queries. To do this, we enlist the LSH Forest self-tuning indexing scheme [37]. This indexing scheme, given a query, will give a subset of nearest-neighbor candidates to which the query can be compared, based on the number of hash collisions between query and candidates. In our case, querying the index with a MinHash signature from a sequencing read will return candidate window signatures.

As in a traditional LSH index, two parameters must be tuned in order to maximize the occurrence of collisions between a query and its nearest neighbor: *i.* the number of hash functions to encode an item (*K*) and *ii.* the number of hash tables (buckets) to split an item into (*L*). As we have already hashed the signature during MinHashing, we are essentially splitting the signature uint64 values over a series of buckets. Explicitly, for each of the *L* buckets we take *K* uint64 values from the signature (where *K* <= *s*) and hash the values again to a single binary string. The original signature of *s* uint64 values is replaced by a more compact representation of *L* binary strings.

The challenge with traditional LSH indexing schemes was setting appropriate *L* and *K* values. Too small a value for *K* results in an increased false positive rate due to increased collisions of dissimilar query-neighbor pairs, and a large value for *K* lowers the collision chances of similar query-neighbor pairs. Therefore, setting *L* > 1 is needed to maximize the occurrence of collisions between similar query-neighbor pairs. To set appropriate *K* and *L* values, the distance between query and nearest neighbor is needed, but tuning for one query-neighbor pair can reduce performance for other query-neighbor pairs [38]. The LSH Forest indexing scheme addresses the limitations of the traditional LSH indexing by using a unique label of variable length to assign data points to buckets and storing these in a data structure that combines multiple prefix trees, each constructed from hash functions (derived from the MinHash signature).

The LSH Forest data structure is first initialised and tuned using the MinHash signature length and the Jaccard Similarity threshold for reporting query-neighbor matches. To tune the index, multiple combinations of the number of buckets (*L*) and the number of hash functions (*K*) are tested (within the bounds of the signature length) for false positive and false negative rates at the specified Jaccard Similarity. *L* and *K* are then set according to the lowest error rate possible and a set of *L* initial hash tables are then created (Algorithm 2 in Additional File 1).

Once the LSH Forest data structure has been initialised, each signature from the variation graph(s) is added to the initial hash tables. Each signature is split into *L* equally sized chunks of *K* hash functions. The chunks are hashed to a binary string (little-endian ordering) and stored in the corresponding hash table, with each chunk pointing to the graph and window from which they derive. Once all signatures have been added, the initial hash tables are transferred into a set of arrays (buckets) and sorted.

### Alignment

To align sequence reads to a variation graph, our method uses an approximate nearest-neighbor search to seed a read against a region of a graph and then employs a hierarchical local alignment to fully align the read.

#### i. Nearest-neighbor search

For a given sequence read, a MinHash signature is generated as during indexing, using the same values for *k* and *s* etc. The signature is then queried against the LSH Forest data structure containing the indexed variation graphs (Algorithm 3 in Additional File 1). To run a query, the signature is split into *L* equally sized chunks and each chunk is hashed to a binary string using the same hash function as during the LSH Forest population [37]. Each hashed chunk of the query signature is then queried against the corresponding bucket in the LSH Forest. To improve querying efficiency, each bucket is compressed by constructing a prefix tree (PATRICIA trie) [39]. Binary search is used to search the bucket (in ascending order) and return the smallest index of the bucket where the prefix matches the query chunk; the bucket is then iterated over from this index position, returning graph windows held until the prefix no longer matches the query chunk. Once all chunks of the query sequence have been searched, all the windows (graph IDs and start nodes) are collated and stored as seeds for each read. The nearest-neighbor search is then repeated using the reverse complement of the query read.

#### ii. Hierarchical local alignment

For a read that has been seeded one or more times, a hierarchical local alignment process is used to align the read to a graph traversal. We assume that most reads do not feature novel variation, so we first try an exact match alignment. We then try an exact match alignment after shuffling the seed *n* nodes along the graph, followed by a clipped alignment. As soon as the read aligns, no further alignments are tried for that seed. To perform an alignment from a given seed, a recursive depth-first search (DFS) of the seeded graph is performed, beginning at the start node identified during the nearest-neighbor search (Algorithm 4 in Additional File 1). If a match between node and read base is encountered, the position in the read is incremented and the next node in the DFS is checked. When no match is found or the whole read has been iterated over, the DFS of the current traversal is ended.

### Reporting

Reads are classified as ARG-derived if they have successfully aligned to a variation graph. In order to apply our method to resistome profiling, the classified reads must be evaluated to annotate what gene they derive from and if that entire gene is present in a given sample.

To classify a read alignment, a score is calculated according to the parent information of the nodes of the alignment. That is, if a node was derived from gene X and gene Y, the read would receive a point for each parent. Points are tallied, the top parent(s) in the tally is used to classify the read. If the top parent was not present for every node traversed, the read is ambiguous. If multiple parents score the highest then the read has multiple valid classifications (a multi-mapper).

Once reads have been classified, gene annotations are then made. Classified reads are binned according to the graphs to which they align and then reference information is extracted from the corresponding graph(s). The length of the reference is used to perform a simple pileup of classified reads. Length and coverage thresholds are applied to determine if a gene can be reported as present in the resistome profile.

### Our implementation

We have implemented our method as an easy to use program called Graphing Resistance Out Of meTagenomes (GROOT) (Figure 2). GROOT is written in Go (version 1.9) and compiles for a variety of operating systems and architectures.

To create the supplied reference data (used for indexing), ARG sequences were downloaded from the CARD, Resfinder and ARG-annot databases [17–19] (accessed June 2016). Each database was clustered using the VSEARCH cluster_size command and stored as MSA files [40]. The GROOT get command will fetch a specified pre-clustered database, check it and unpack it ready for indexing. Alternatively, a user can provide their own clustered database.

The index command checks MSA files for formatting and discards sequences shorter than the expected read query length. Variation graphs are then built from the MSAs, pruned and topologically sorted. For each graph, a sliding window is moved along each traversal (default length = 100, offset = 1), decomposed to k-mer sets (default k=7) and MinHash signatures are created using the Go implementations of Spooky and Farm hash functions (default signature length = 128, based on XORing the Spooky and Farm hash functions) [41–44]. An LSH Forest is then initialised, tuned and populated, before being serialised using the Go gob package and written to disk.

The align command loads the index, sets all hashing parameters to match the index and then streams the FASTQ read file(s). MinHash signatures are created for each read and its reverse complement; signatures are then queried against the index. Once seeded, reads are optionally quality trimmed prior to hierarchical local alignment and reporting.

The index, align and report subcommands of GROOT all utilise a concurrent pipeline pattern that is driven by the flow of data between structs. This pattern also facilitates the streaming of data from STDIN, as well as from disk, and allows the GROOT commands to be piped together.

### Evaluating performance

The full commands and code used to evaluate the performance of our implementation can be found in the GROOT repository (https://github.com/will-rowe/groot/paper). For running the accuracy benchmark, simulated FASTQ reads (150bp read length) were generated from the ARG-annot database using BBMap [19, 45]. An index of the ARG-annot database was created using the GROOT index command with default settings. Reads were aligned using the GROOT align command with default settings, running on a Linux laptop using 1, 4, and 8 CPUs for each test respectively.

For running the comparison benchmark, the genomes from the Critical Assessment of Metagenome Interpretation (CAMI) project were downloaded in FASTA format [46]. The genomes were screened against the CARD database to mask any ARGs already present in the genomes [18]. A subset of ARGs were then randomly sampled from the CARD database (v1.1.2) using Bioawk and were combined with the CAMI genomes in a single FASTA file [18]. The CARD database was selected as ARGs-OAP uses this and it is one of the default databases provided by GROOT [23]. Sets of metagenomic FASTQ reads (150bp read length, errors allowed) at varying coverage levels were then generated from the FASTA file using BBMap [45]. For each set of metagenomics reads, GROOT and ARGs-OAP (stage one) were both run with default parameters on a LINUX laptop using 1 core [23]. ARGs-OAP (stage two, version 2) was completed on the dedicated Galaxy webserver, allowing only full-length, 100% identity read matches [23].

## Results

The results presented here evaluate our implementation of indexed variation graphs for resistome profiling, in terms of both the accuracy of the tool and its performance compared to a recently published resistome profiling tool.

### Accuracy assessment

To assess the accuracy of our implementation, we generated sets of random FASTQ reads from the ARG-annot database, classified the reads with GROOT and then compared the classification to the ARG from which the read derived [19]. For six sets of simulated reads, ranging from 100 to 10,000,000 reads, GROOT classified the ARG-derived reads with a false negative rate of 0.000001%. GROOT took an average of 299.53 wall clock seconds to classify 1000000 ARG-derived reads using 8 CPUs. At <1000 reads, runtimes were equivalent regardless of the number of CPUs used, however as the datasets approached real-world metagenome size (>1,000,000 reads), increasing the number of CPUs resulted in decreased runtime of approximately 2.5-fold per 4 CPUs (Figure 3).

**Figure 3.**
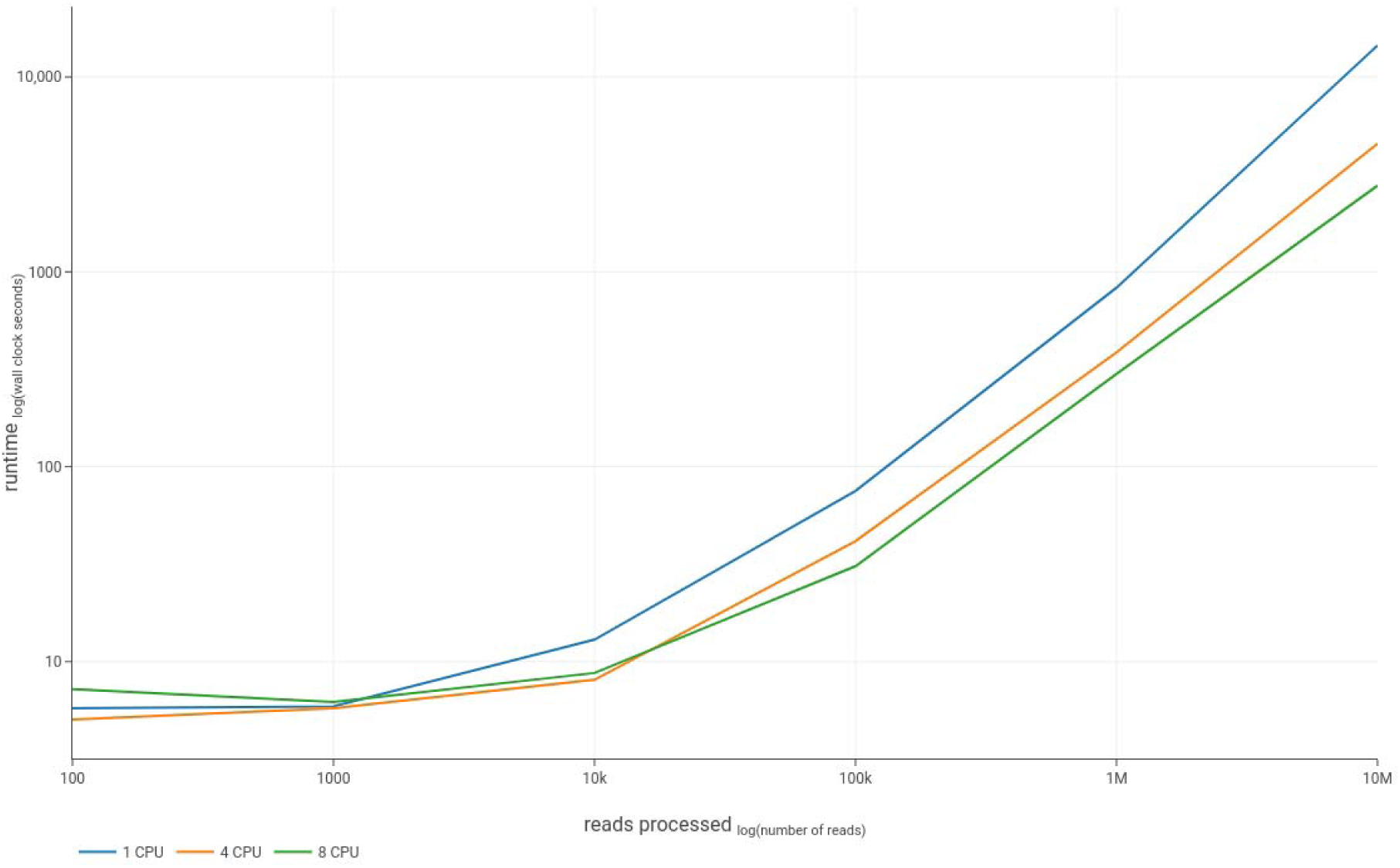
Runtime performance. This figure shows the runtime performance of our implementation (GROOT) on 1, 4, and 8 CPUs. The data was collected during the accuracy assessment of GROOT, where it classified ARG-derived read with a false negative rate of 0.000001%.

### Performance comparison

To compare the accuracy and speed of our implementation against a recently published resistome profiling tool (ARGs-OAP [23]) we used a set of publically available, simulated metagenomes and spiked in full-length ARG sequences. The spiked data was sampled at varying coverage levels and the performance of GROOT and ARGs-OAP was evaluated in terms of the number of ARGs correctly annotated from each sample, as well as the time taken for each tool to process the samples.

On average GROOT was 6.3 times faster at processing the samples compared to stage one of ARGs-OAP. The mean time to process a metagenome with 1X coverage (approximately 1.1 million reads) was 157.87 seconds for GROOT, compared to 989.15 seconds for ARGs-OAP (stage one only) (Figure 4.a). ARGs-OAP stage two took an average of 12 hours to run on a dedicated server (not included in Figure 4.a.).

**Figure 4.**
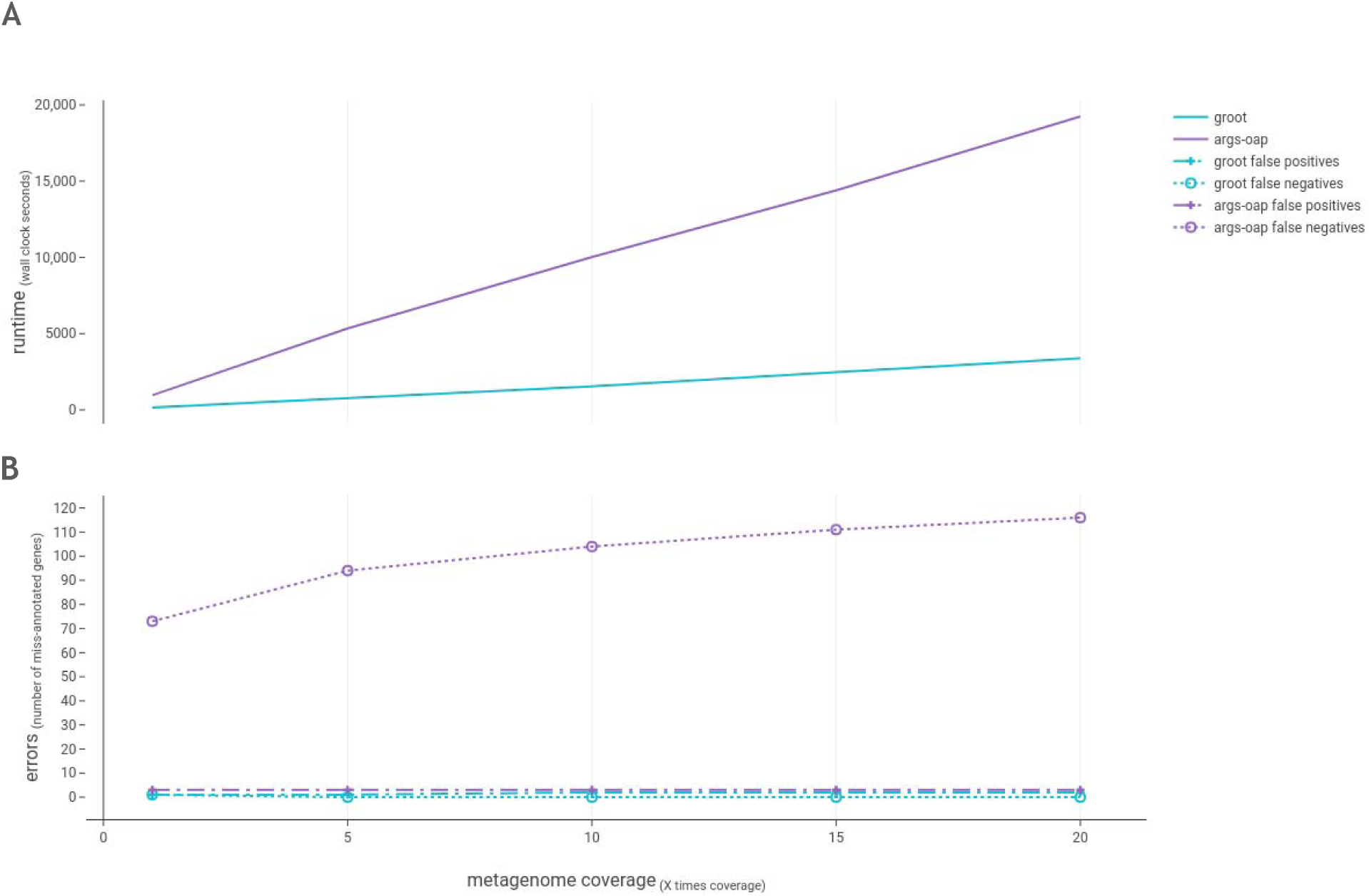
Benchmarking GROOT and ARGs-OAP. Figure 4.a shows the runtime comparison of GROOT and ARGs-OAP when processing metagenomes at varying coverage levels. Figure 4.b shows the number of errors (false positives and false negatives) recorded by each program at varying metagenome coverage levels.

In terms of accuracy, GROOT recorded only one false negative across all samples (present in the lowest coverage metagenome), whereas ARGs-OAP (Stages 1 + 2) consistently recorded three false negatives per sample. GROOT had a mean rate of 1.6 false positives per sample, whereas ARGs-OAP had a mean rate of 99.6 false positives per sample. The number of false positives found by GROOT did not increase beyond 10X coverage, compared to the false positive increase observed at every coverage increase by ARGs-OAP (Figure 4.b).

## Discussion

Antimicrobial resistance remains a significant challenge to human and animal health. With the increasing drive for AMR surveillance in order to inform policy and mitigate the spread of resistance, metagenomic studies to identify and monitor ARGs in a wide variety of biomes are becoming commonplace. Sampled biomes range from the human gut, through to water supplies, agricultural land and marine environments. Despite this scale and variety, the question being asked remains simple, what ARGs are present in a given sample? Although simple, resistome profiling still presents a challenge, as illustrated by the many tools available (Figure 1). Our method improves upon two main issues with these existing tools; the resolution offered and the speed at which a sample can be profiled. Crucially also, as a novel algorithm, it is translatable to identifying other highly-related gene sets within metagenomes.

In terms of profiling resolution, we are referring to the accuracy of the annotations in a resistome profile. That is, is it possible to annotate the sub-type of ARG in a sample (e.g. *bla-SHV-1*) or just the type (e.g. *bla-SHV*)? This resolution is important as different ARG subtypes can provide selective resistance to different antibiotics, for example genes within a single beta lactamase gene class (structural classes, based on amino acid sequence) can confer resistance to different beta lactam antibiotics [25]. Therefore, the greater the resolution, the more information can be extracted from a sample. To gain this resolution, we need to both cover an entire reference sequence and be confident in the reads placed. We address this by allowing only exact matches to the whole reference sequence before annotating the gene as present. The counter to this argument for greater resolution is that sequence quality will impact read placement and also, novel ARGs will be missed. Whilst we believe that the main utility of GROOT is its ability to confidently resolve ARGs, we have also added features to allow for these points. Firstly, our implementation has an optional quality trimming algorithm to remove low quality bases prior to read alignment. Secondly, to allow potentially novel ARGs or accommodate low coverage samples, there is an option to relax the scoring system in order to allow non-exact matches or partially covered genes.

Our method offers an improvement in resolution over those that consolidate variant ARGs to a representative sequence [14, 20], or that have high false positive rates due to allowing partial or inexact matches [23]. Despite a marked improvement on ARGs-OAP in our benchmark our method did record a false negative and some false positives, likely as a result of introduced sequencing error and the low sample coverage (in the case of the single false negative). These errors could be considered a limitation of the exact local alignment utilised in this method, a more relaxed alignment could be allowed but this would be at the expense of confidence in the ARG annotations.

In terms of speed, our method offers several advantages over other resistome profiling tools. Our method does not require metagenome assembly or the upload of data to remote servers, both of which add significant time to a resistome profiling analysis. The latter requires good bandwidth and the former can require large compute resources; a complex metagenome can take up to 10 hours and 500GB of RAM to assemble [47]. Our implementation can run a typical 2 gigabyte metagenome in 2 minutes using a single CPU, and scales when run on higher-performance systems. Our benchmark was restricted to a single CPU as ARGs-OAP is limited in the number of CPUs it can use. The benchmark also ignored the time taken during stage 2 of ARGs-OAP (on the remote server) as this could be variable depending on server load and available bandwidth for upload. Despite this, GROOT still offers a much faster runtime than ARGs-OAP. In addition to runtime, analysis times are also reduced with GROOT due to its ease of use. It runs as a self-contained binary, is packaged with bioconda [48] and requires only two commands to run a resistome profiling analysis, offering significant advantage over more complex workflows or those that require upload to remote servers halfway through the analysis [23]. Our implementation is targeted towards researchers who may not have access to high performance computing and wish to run metagenomics workflows on a laptop. However, GROOT also scales for use on larger servers.

With the advent of long-read and barcoding protocols for resistome profiling, it is tempting to think that tools for short read resistome profiling may prove less relevant in the near future [49, 50]. However, Illumina sequencing currently remains the standard technology for metagenomics applications and historic Illumina datasets will also need to be reanalyzed where new sampling is not feasible. In addition, our method can be used in conjunction with other new methods that elucidate gene context in metagenomics samples, allowing for novel insights to be gained from analysis and re-analysis of metagenomic data collections [51].

Our implementation offers efficient and accurate resistome profiling, enabling the identification of full-length ARGs in complex samples. It should be noted that this method is database dependent and is not intended for the identification of novel ARGs. The implementation is easy and quick to run as there is no complicated installation or dependencies, no split local/remote processing, it facilitates streaming of input and is built using Go’s concurrency patterns.

## Conclusions

We present a method for resistome profiling that utilises a novel index and search strategy to accurately type resistance genes in metagenomic samples. The use of variation graphs yields several advantages over other methods using linear reference sequences. GROOT performed much more quickly than a recent resistome profiling tool, and also recorded few false positives and negatives. Our method is not restricted to resistome profiling and has the potential to improve current metagenomic workflows.

## List of abbreviations

AMR: Antimicrobial Resistance
ARG: Antimicrobial Resistance Gene
DAG: Directed Acyclical Graph
DFS: Depth-First Search
GROOT: Graphing Resistance Out Of meTagenomes
MSA: Multiple Sequence Alignment
PCR: Polymerase Chain Reaction

## Declarations

### Availability of data and materials

The source code for our implementation, as well as the code used to evaluate its performance and plot the manuscript figures, can be found in the GROOT repository (https://github.com/will-rowe/groot) (MIT License. DOI: https://doi.org/10.5281/zenodo.1186232).

### Author contributions

WPMR conceived and implemented the method. All authors wrote, read and approved the final manuscript.

### Ethics approval and consent to participate

Not applicable.

### Consent for publication

Not applicable.

